# Graph Convolutional Network-based Method for Clustering Single-cell RNA-seq Data

**DOI:** 10.1101/2020.09.02.278804

**Authors:** Yuansong Zeng, Jinxing Lin, Xiang Zhou, Yutong Lu, Yuedong Yang

## Abstract

Single-cell RNA sequencing (scRNA-seq) technologies promise to characterize the transcriptome of genes at cellular resolution, which shed light on unfolding cell heterogeneity and diversity. Fast-growing scRNA-seq profiles require efficient clustering algorithms to identify the same type of cells. Although many methods have been developed for cell clustering, existing clustering methods are limited to extract the representations from expression data of individual cells, while ignoring the high-order structural relations between cells. Here, we proposed GraphSCC, a robust graph artificial intelligence model to cluster single cells by accounting for structural relations between cells. The representation learned from the graph convolutional network, together with another representation output from a denoising autoencoder network, are optimized by a dual self-supervised module for better cell clustering. The experimental results indicate that GraphSCC model outperforms state-of-the-art methods in terms of various evaluation metrics on both simulated and real datasets. Further visualizations show that GraphSCC provides representations for better intra-cluster compactness and inter-cluster separability.

## I. Introduction

Single-cell analysis is a valuable tool for discovering cellular heterogeneity in complex tissues and diseases[1, 2]. Clustering is an essential step in single-cell analysis, since each cell cluster represents a distinct cell state or type in transcriptome space. Despite the significant improvements in measuring scRNA-seq technologies and advances of many clustering methods, it remains challenging for clustering cells based on scRNA-seq data [3]. Concretely, scRNA-seq data often contains dropout events and substantial noise due to biological and experimentally technical factors, such as amplification bias, the low RNA capture rate [4], and cell cycle effects [5]. A dropout event is defined as missed gene measurements, resulting in a ‘false’ zero count observation [6]. Thus, solving the dropout events and substantial noises is important for improving clustering analyses.

Several imputation methods have been developed for solving the dropout events of scRNA-seq data. Early methods are often based on statistical models, e.g., CIDR [7], scImpute [8], MAGIC [9], and SAVER [10]. Due to deep learning techniques achieving state-of-art results in many areas, several researchers developed neural-network-based imputation methods. For example, DCA [6] reconstructs the scRNA-seq data through the autoencoder optimized by a loss function of the zero-inflated negative binomial (ZINB) [11]. DeepImpute uses highly correlated genes and sufficient reads coverage to recovery missing values[12]. GraphSCI employed graphical neural network to capture the relations between genes for accurate imputations [13]. Although the imputed scRNA-seq data help improve the clustering results, the results remain unsatisfactory. These imputation methods are not optimized for cell clustering, and the imputed data by imputation methods may produce false-positive gene-gene correlations.[14].

Recently, a few clustering methods have been specifically designed for scRNA-seq data. For example, the spectral clustering method SIMLR learns a robust distance metric to fit the structure of scRNA-seq data[15]. Seurat3.0 applies the Louvain algorithm [16] to cluster cells depended on the low-dimensional scRNA-seq data [17]. DendroSplit through feature selection in scRNA-seq data to uncover multiple levels of biologically meaningful cell populations [18]. ScDeepCluster is a deep learning embedded clustering method, which accounts for the overdispersion and sparsity of the scRNA-seq when clustering [19]. There are a few tools had been developed for dividing single cells into hierarchies or groups, such as SC3[20], RaceID[21], SNN-Cliq[22], BISCUIT[23], and pcaReduce[24]. However, most of these methods rely on only the data of individual cells without explicitly considering structural relations between cells.

The Graph Convolutional Networks (GCN) can efficiently capture structural information[25]. In recent years, GCN and its variants [26, 27] have been successfully applied to a wide range of applications, including protein prediction [28], traffic prediction [29] and drug design[30]. Xie et al. developed a deep embedding method for clustering analysis (DEC)[31] by unsupervised manner, which uses an auxiliary target distribution to iteratively refines clusters by learning highly confident assignments. DEC and its variant IDEC[32] have been successfully used in molecular biology[19, 33]. Recently, Bo et al. developed a Structural Deep Clustering Network (SDCN) for integrating structural information between objects[34]. Theoretically, they have proved that the inclusion of GCN enables a high-order regularization constraint to learn better representations that help improve the clustering results, and SDCN outperformed other methods in many types of datasets.

Inspired by these works, we present a robust graph-based artificial intelligence model, GraphSCC, to integrate structural information in the clustering of scRNA-seq data. Meanwhile, we employed a denoising autoencoder network to obtain low dimensional representations for capturing local structural. A dual self-supervised module was then employed to optimize the representations and the clustering objective function iteratively in an unsupervised manner. The results show that the GraphSCC outperforms state-of-the-art methods on both real datasets and simulated. Furthermore, GraphSCC provides representations for better intra-cluster compactness and inter-cluster separability in the 2D visualization.

The advantages of GCN is its native learnable properties of aggregating and propagating attributes to obtain relations over the whole cell-cell graph. Thus, the learned graph representations can be treated as high-order representations between cells. The superior performance of GraphSCC in cell cluster prediction benefits from *(i)* we synergistically determine cell clusters based on the integration of high-order topological relations between cells and characteristics of individual cells, and *(ii)* we apply the dual self-supervised module to iteratively refine clusters by learning from highly confident assignments using an auxiliary target distribution.

## II. Materials and Methods

### A. Datasets and Preprocessing

#### Simulated Data

We applied a generally used R package Splatter to generate simulated scRNA-seq count data [35]. For all simulated data, we set 2000 cells composed of 2000 genes with four groups of the same numbers, i.e., 500 cells per group. Following the previous study[19], we set *dropout. mid* = 2, *dropout. shape*= −1(fixed dropout rates at 45%), and used default values for other parameters. To simulate various clustering signal strengths, we generated datasets with different de.fracScale in {0.2, 0.225, 0.25, 0.275, 0.3, 0.325, 0.35, 0.4}. The de.fracScale is the parameter sigma of a log-normal distribution to control multiplicative differential expression factors. To avoid random fluctuations, we repeatedly generated 20 datasets for each setting with different random seeds, and reported the average results.

#### Real Data

We downloaded 15 datasets of human and mouse scRNA-seq involved in various tissues and different biological processes as used in the previous study[36] from the Hemberg group (https://hemberg-lab.github.io/scRNA.seq.datasets/). The datasets contain different scales of cells from dozens to thousands derived from various single-cell RNA-seq techniques. The detail information of datasets was listed in TABLE I. The data type of top 10 datasets is raw read counts, and the last 5 datasets are normalized counts.

**TABLE I.** THE LIST OF DTATSETS USED IN THIS STUDY.

| Datasets | GSE/ID | #Cells | #Genes | #Cell types |
| --- | --- | --- | --- | --- |
| Baron Human | GSE84133 | 8569 | 20125 | 14 |
| Baron Mouse | GSE84133 | 1886 | 14878 | 13 |
| Darmanis | GSE67835 | 466 | 22088 | 9 |
| Deng | GSE45719 | 268 | 22431 | 6 |
| Goolam | E-MTAB-3321 | 124 | 41427 | 5 |
| Klein | GSE65525 | 2717 | 24175 | 4 |
| Li | GSE81861 | 561 | 55186 | 9 |
| Romanov | GSE74672 | 2881 | 24341 | 7 |
| Segerstolpe | E-MTAB-5061 | 3514 | 25525 | 15 |
| Zeisel | GSE60361 | 3005 | 19972 | 9 |
| Biase | GSE57249 | 56 | 25733 | 4 |
| Tasic | GSE71585 | 1679 | 24150 | 18 |
| Treutlein | GSE52583 | 80 | 23271 | 5 |
| Xin | GSE81608 | 1600 | 39851 | 8 |
| Yan | GSE36552 | 90 | 20214 | 6 |

#### Preprocessing

We normalized simulated scRNA-seq counts using the transcripts per million (TPM) method [37] and then scaled the value of each gene to [0, 1]. For real datasets, we followed Seurat3.0’s procedure to normalize and select the top 2000 highly variable genes for scRNA-seq data and then scale each gene’s value to [0,1]. Note that for real datasets normalized by FPKM, we first converted them to TPM by Eq. (1) as proposed by [38], and then preprocess the data as above.

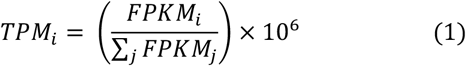

### B. GraphSCC Architecture

GraphSCC network consists of three components as shown in Fig. 1: Denoising Autoencoder (DAE), Graph Convolutional Network (GCN), and Dual Self-supervised Module (DSM). DAE network learns robust low-dimensional representations that could reconstruct the inputs. GCN optimized the representations with constraints from structural information between cells. DSM applied to cluster the cells according to the learned representations by DAE and GCN.

**Fig. 1.**
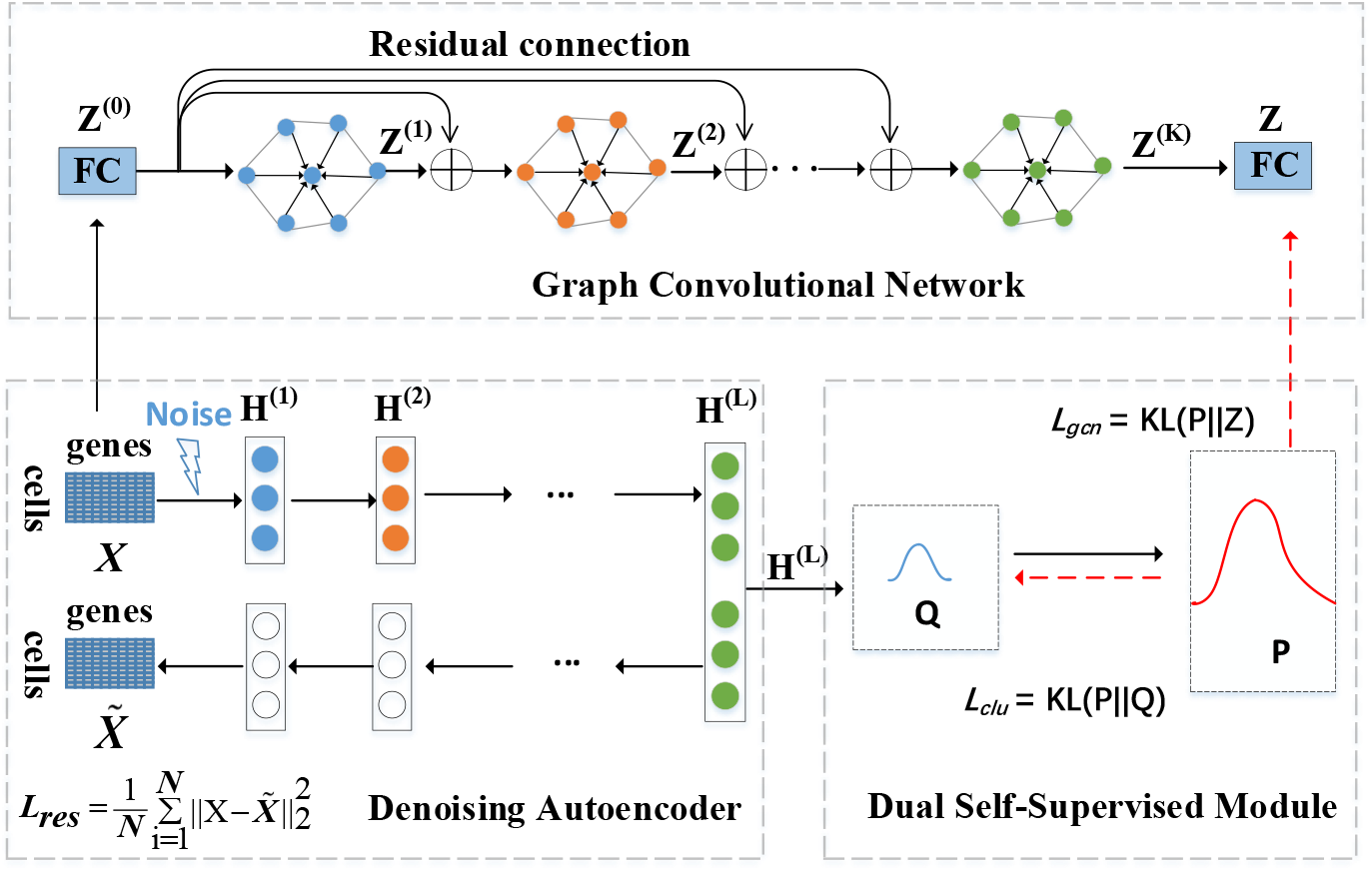
The framework of our proposed GraphSCC consisting of the Denoising Autoencoder (DAE), Graph Convolutional Network (GCN), and Dual Self-Supervised Module (DSM). The ⊕ is the residual connection block on the initial representation *Z*^(0)^.

#### 1. DAE Networks

The gene expression is represented by the matrix *X* ∈ ℝ^*N*×*d*^, where *N* is the number of single-cell and *d* is the dimension of expressed genes. DAE encodes gene expression matrix to fixed-size vector representations for reserving local structure. DAE is a variant version of autoencoder that is input with corrupted data and outputs the fit data. Concretely, at encoding layer *𝓁* the output *H*^(*𝓁*)^ is computed as:

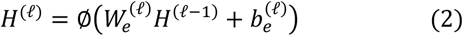

where ∅ is the activation function, 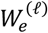 and 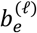 are the weight matrix and bias parameters, respectively. *H*^(0)^ = *X* + *ϵ*, and *ϵ* is the Gaussian noise. The decoded layers are caculated as:

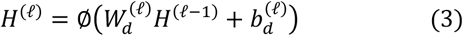

where ∅ is the activation function, 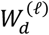 and 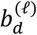 are the weight matrix and bias parameters, respectively,and the output of the last layer is the reconstructed data 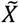. The encoder and decoder networks are both fully connected neural networks using the RELU activation function. The weight and bias parameters are optimized using the following loss function:

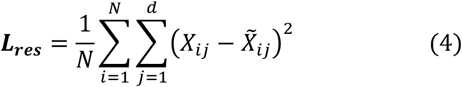

#### 2. KNN Graph

The GCN graph was constructed by KNN, where the cell similarity is calculated by the dot product function as:

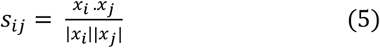

where *x*_*i*_ and *x*_*j*_ are the embedded features for cells *i* and *j*, the embedded features *x*_*i*_ and *x*_*j*_ obtained from the representation *H*^(*L*)^ of pre-train DAE, and |*x*_*i*_| *and* |*x*_*j*_| are the corresponding modules, respectively. According to the similarity matrix *S* ∈ ℝ^*N* ×*N*^, the most similar cells of every cell are selected as its neighbors to construct the adjacency matrix *A* for GCN. For all datasets, the number of each cell’s neighbors was at most 1% of the total number of cells with a maximum of 20.

#### 3. GCN Network

We apply the GCN network to capture structural information between cells that the DAE network has ignored. Inspired by a recent study [39]. We alleviate the well-known over-smoothing phenomenon in GCN using the residual connection [25]. Since the large feature dimension d of *X*, a lower-dimensional *Z*^(0)^ used as the initial representation of GCN, which is extracted from input feature *X* using a fully-connected neural network as follows:

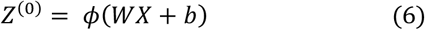

where *Z*^(0)^ ∈ ℝ^*N* ×*m*^, as the initial representation of GCN, m is lower than d. *W* and *b* are weight matrix and bias parameters, respectively.

Then, the representation *Z*^(*k*+1)^ learned by GCN can be obtained by the following convolutional operation:

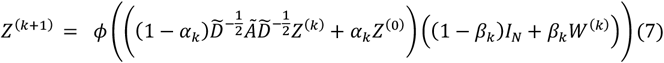

where the hypermeter *α*_*k*_ > 0 ensures each layer retaining the information from input layer *Z*^(0)^ and the hypermeter *β*_*k*_ > 0 ensures the decay of the weight matrix adaptively with stacked layers. *Ã* = *A* + *I*_*N*_ with A as the adjacency matrix obtained from the KNN-graph and *I*_*N*_ is the identity matrix. 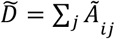 is the normalization term. The *ϕ* is the RELU activation function. We set 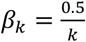 and *α*_*k*_ = 0.3 following the previous study [39].

The last layer in GCN module is connected using a *softmax* function:

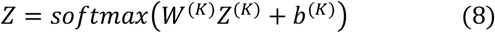

where the output Z∈ ℝ^*N*×*c*^ could be treated as the probability distribution, and *c* is the number of cell clusters. The result *z*_*ij*_ ∈ *Z* means the probability cell *i* belongs to cluster center *j*.

#### 4. Dual Self-supervised Module

We designed the dual self-supervised module to help DAE and GCN learn low-dimensional representations for better cell clustering. Through the input of *H*^(*L*)^ from DAE, the cells are grouped into *c* clusters corresponding with *c* cluster centers *u*_*j*_, *j* = 1, …, *c* through the *K*-means algorithm. With the help of an auxiliary target distribution, the clusters are refined iteratively until convergence by learning from high confidence assignments.

Specifically, for the *i*-th single cell and the *j*-th cluster, the Student’s t-distribution [40] is applied as the kernel to compute the similarity between the cluster center vector *u*_*j*_ and the data representation *h*_*i*_ as follows:

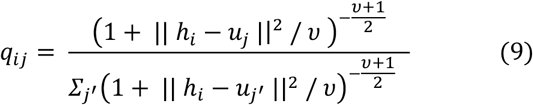

where *v* is the degree of freedom of the Student’s t-distribution (*v* set as 1 in this study), and *q*_*ij*_ can be regarded as the probability of *i*-th cell belonging to *j*-th cluster. Based on the calculated distribution *Q* = [*q*_*ij*_], the target distribution *P* = [*p*_*ij*_] is computed by:

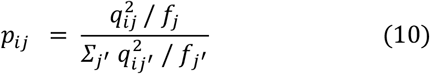

The procedure of GraphSCC is also shown in Algorithm 1.

#### Algorithm 1: Training process of GraphSCC

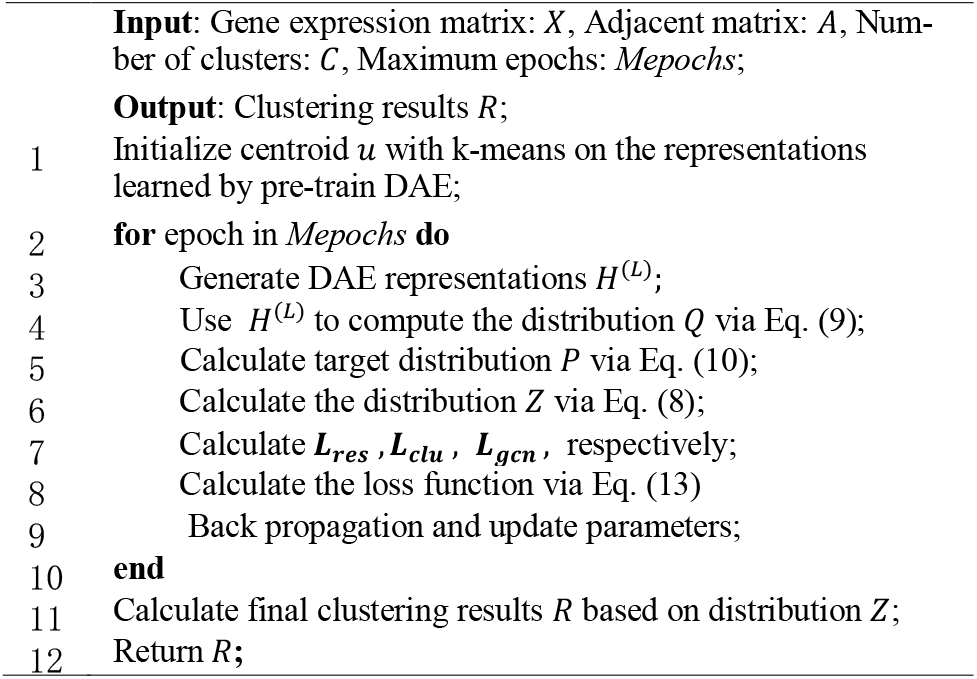

where *f*_*j*_ = *∑*_*i*_ *q*_*ij*_ is soft frequency for cluster *j*. To better clustering, the loss function could be defined to minimize the Kull-back-Leibler (KL) divergence between two probability distributions as:

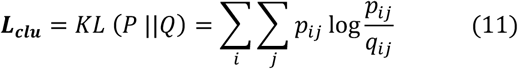

By minimizing the loss *L*_*clu*_ between the distribution *Q* and *P*, the target distribution *P* helps the DAE module learn better low-dimensional representations for clustering cells. Since the target distribution *P* is defined by *Q*, minimizing *L*_*clu*_ is a form of self-supervised learning mechanism.

To combine the structural information between cell-to-cell, the target distribution *P* was applied to supervise the updating of distribution *Z* as follows:

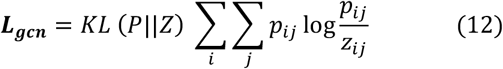

By optimizing the distance between the target distribution *P* and soft assignments *Z*, the GCN module’s parameters will learn useful information from the DAE module. In this way, the GCN representations contain both the structural information and the characteristic information of data. Since the target distribution *P* supervises the DAE and GCN modules simultaneously, we call it a dual self-supervised mechanism.

Finally, the total loss function of GraphSCC is defined as:

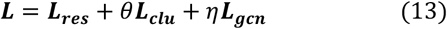

where *η* and *θ* are two hyper-parameters to balance contributions from structure information of scRNA-seq data and the clustering optimization. We set *θ* = 0.1 and *η* = 0.01 for all the datasets.

Since the GCN’s representations contain structural information and characteristic information, the distribution *Z* is used as the final clustering results. Thus, the label assigned to cell *i* is:

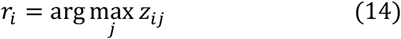

where *z*_*ij*_ is computed in Eq. (8) for cell *i* in the *j-*cluster.

### C. Training and Evaluation

#### 1. Evaluation Metrics

The clustering results are evaluated by three commonly used metrics, Clustering Accuracy (CA) [41], Normalized Mutual Information (NMI) [42], and Adjusted Rand Index (ARI) [43]. The NMI is defined as:

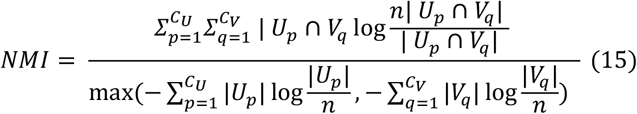

where *U* and *V* are the predicted and truth assignments of totally *n* cells into *C*_*V*_ and *C*_*U*_ clusters, respectively. The numerator is the mutual information between *U* and *V*, and the denominator is the entropy of the clustering *U* and *V*.

The CA is explained as the best match between the predicted cluster assignments and the truth assignments, calculated as:

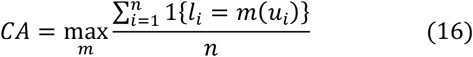

Where *n* is the number of data points, and *m* ranges over all probable one-to-one mapping between real label *l*_*i*_ and clustering assignment *u*_*i*_.

The ARI is defined as:

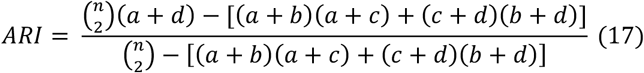

where *a* is the number of cell pairs belonging to the same group in both *U* and *V, b* is the number of cell pairs belonging to different groups in *V* and the same group in *U, c* is the number of cell pairs belonging to the same group in *V* and different groups in *U*, and d is the number of cell pairs belonging to different groups in *U* and different groups in *V*.

#### 2. Implementation

The GraphSCC model was implemented in python 3 using PyTorch. The dimensions of DAE is set to d-512-256-64-10, where d is the dimension of the input data, and 10 is the dimension of the bottleneck latent *H*^(*L*)^. DAE is first pre-trained for 400 epochs by the optimizer Adam with the initial learning rate *lr* = 0.0001 and the batch size of the pre-train equaling to 32. We used the “Randn” function in PyTorch to generate Gaussian noises. Note that, for simulated data, we reduce the noise value to 0.2 times its value. We set the layers of GCN as 5, and set the dimensions of the initial representation *Z*^(0)^ and the hidden layer of GCN as 256. The optimizer for the clustering stage is Adam with setting *lr* = 0.00001, and the clustering training epoch is 1000. The training stops until the proportions of cells to change clustering assignment (*ca*) are below a threshold *tol* in 300 consecutive steps. The *ca* is computed as 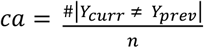, where *n* is the number of all cells, *Y*_*curr*_ is the cluster identity gained by the maximum cluster assignment possibility in the current step, *Y*_*prev*_ is the corresponding identity in the previous step, and *#*|*Y*_*curr*_ *≠ Y*_*prev*_ | is the number of cells whose *Y*_*curr*_ differ from *Y*_*prev*_. We set *tol* = 0.0001 by default. For all other competing methods, we use the default parameters provided in the original articles. All experiments were conducted on the Nvidia Tesla P100 (16G).

#### 3. Hyper-parameters tunning

There are multiple hyperparameters in our model. Here, we tested essential hyperparameters and the scope of values as follows:

1. **DAE module:** The larger bottleneck layer size of DAE may explain more variations. Thus, we tested {2, 5, 10, 20, 32, 64} for the size of the middle layer and found 10 is the optimal value. The pre-training epochs may affect the clustering centroid initialized by K-means. We tested the following settings {50, 100, 200, 400, 600} and found 400 epochs to be the optimal value.
2. **GCN module:** The higher number of GCN layers means the deeper information aggregated from the node and edge features, while excessive layers may result in tremendous computing resources and vanishing gradients. Here, we tested the following settings {2, 3, 4, 5, 8, 16} and found 5 layers to be the optimal value.
3. **Dual self-unsupervised module:** The training epochs may affect the convergence of the model. Thus, we tested the training epochs range from epochs 400 to 1500 with a step of 100 and found 1000 epochs can achieve the best performance.

## III. Results

### A. Evaluation of GraphSCC

To investigate the performance of GraphSCC in different sceneries, we employed R package Splatter to generate simulated scRNA-seq data with different values of the parameter sigma. A greater sigma value means more significant distances between cells from different cell clusters with lower clustering difficulty. As shown in Fig. 2, GraphSCC consistently outperformed the competing methods for NMI values. Though methods Seurat3.0 and IDEC can reach the same NMI value (∼0.98) as GraphSCC at sigma of 0.4, Seurat3.0 and IDEC have a sharp drop to 0.25 and 0.37, respectively at sigma of 0.2. In contrast, GraphSCC is much flatter with an NMI value of 0.76 at a sigma of 0.2. Methods CIDR, scDeepCluster, and DCA performed badly at the sigma of 0.2 with an NMI value below 0.1, but they can achieve decent results at the sigma of 0.4 with NMI values of 0.72, 0.77, and 0.9, respectively. The SIMLR failed to explore clustering signals, which is consistent with the previous observation [19]. Similar trends could be observed for CA and ARI values **(Figure S1)**.

**Fig. 2.**
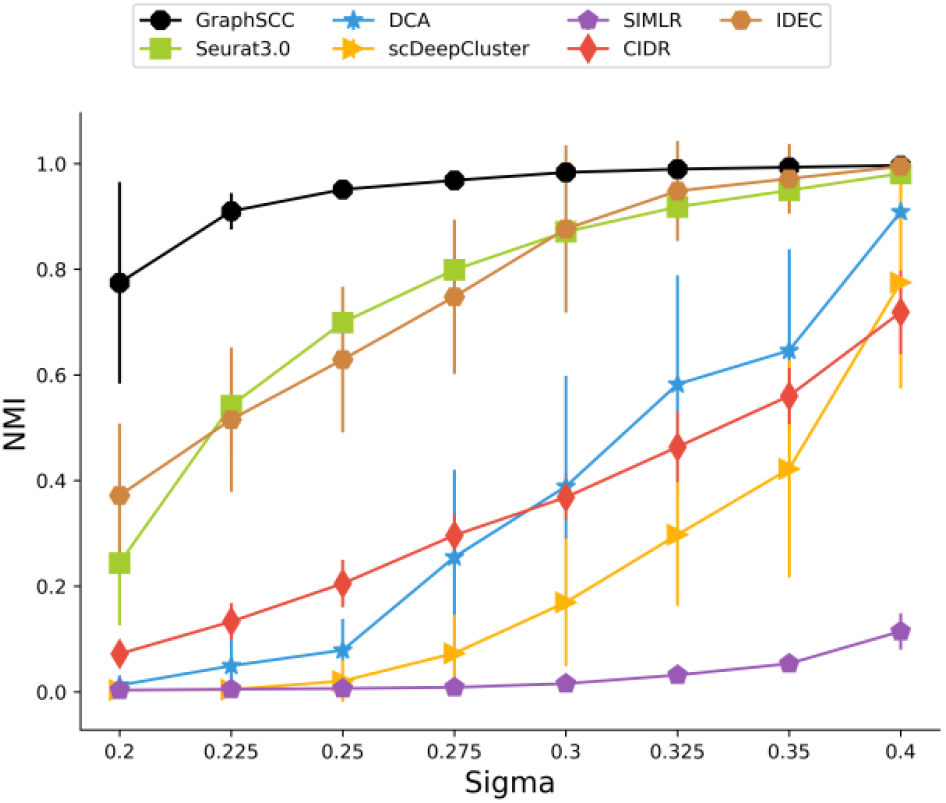
The average and mean square error values of NMI on 20 groups of simulated scRNA-seq data at different sigma values. A higher sigma value represents a stronger signal, corresponding to easier datasets.

We further evaluated GraphSCC on real scRNA-seq datasets with different species and tissues (Details in TABLE I). As shown in Fig. 3, GraphSCC had superior cell clustering results compared to other competing methods on all evaluation metrics. On average, GraphSCC achieved 0.798, 0.791, and 0.744 for the CA, NMI, and ARI, respectively (**Detail information seen in Table S1-3**). These are respectively 13.3%, 9%, and 19% higher than those achieved by the 2nd best method Seurat3.0. Methods scDeepCluster and SIMLR achieved comparable NMI to Seurat3.0 and ranked the 4th and 5th. CIDR has an average NMI value of 0.64. IDEC obtained the lowest NMI value. These methods had generally similar ranks when measured by CA or ARI. The consistent superior results of GraphSCC over the simulated and real datasets demonstrated the robustness of our method.

**Fig. 3.**
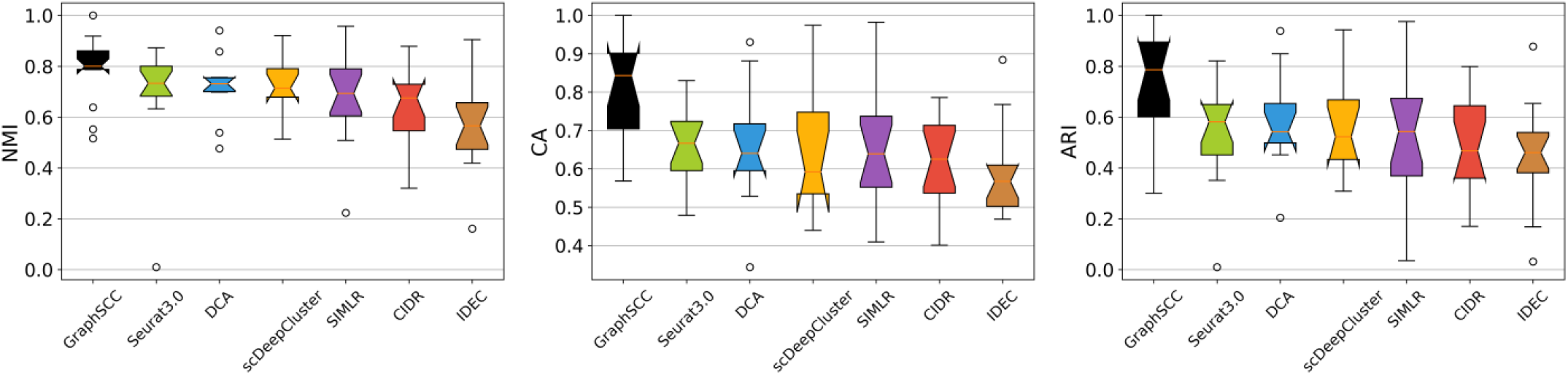
The boxplots of CA, NMI and ARI values for each clustering method on 15 real datasets.

### B. Illustration of the GraphSCC

In this section, we visualized the clustering results on three datasets with different scale of cells. To illustrate the embedded representation effectiveness of GraphSCC, we employed t-SNE[40] to visualize embedded representation in the two-dimensional (2D) space. As shown in Fig. 4. On the Baron Mouse dataset, DCA, CIDR and scDeepCluster showed poor performances. Seurat3.0 and SIMLR showed better separation, but the beta cells (colored blue) were separated into at least three groups and mixed with alpha cells. Compared to Seurat3.0, GraphSCC separated beta cells in one group and produced more compact clusters for all cell types. Alpha and gamma cells were mixed both in GraphSCC and Seurat3.0. Similar trends could be observed for other methods **(Figure S2-3)**.

**Fig. 4.**
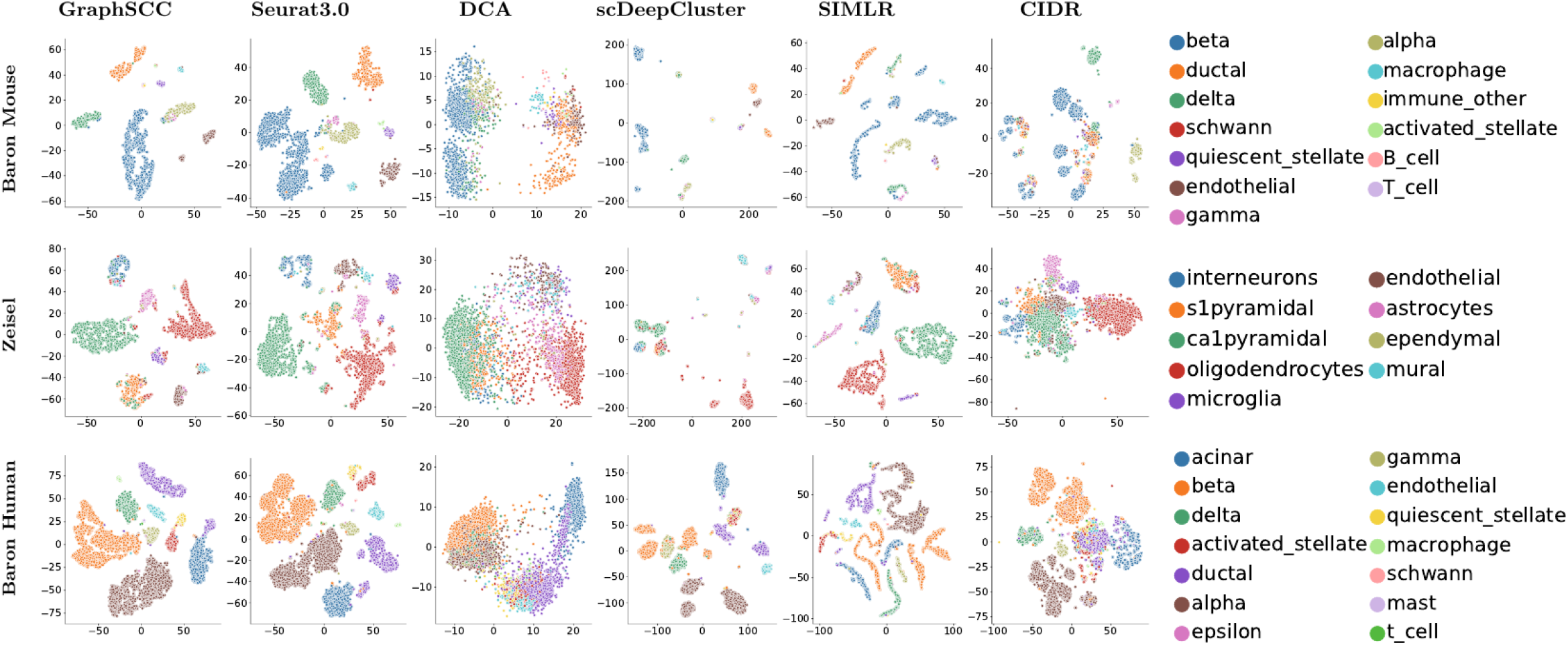
Comparison of the embedded representations of each method in 2D space using the visualize method t-SNE with each point representing a cell and colors for the true cell labels.

On the Zeisel dataset, DCA, scDeepCluster, and CIDR showed poor performances in classification, which mixed a few clusters together. Seurat3.0, SIMLR, and GraphSCC separated most cell populations. In comparison, the cell clusters in GraphSCC and SIMLR have better intra-cluster compactness and inter-cluster separability than Seurat3.0, especially in the oligodendrocytes and ca1pyramidal cells. All methods failed in separating the ca1pyramidal and s1pyramidal cells.

On the Baron Human dataset, DCA performed bad and only separated three groups. For SIMLR, CIDR, and scDeepCluster, at least seven compact clusters of cells were separated, but they were underclustering in the alpha cells (colored brown), where the cell types were separated into at least three groups. In contrast, Seurat3.0 and GraphSCC separated the most cell populations, while Seurat3.0 mixed a few beta and ductal cells with the alpha cells. All methods including GraphSCC separated the ductal cells into at least two groups.

We further visualized the wrongly clustered cells by the Sankey river plots on the Baron Mouse dataset as shown in Fig. 5, where GraphSCC achieved CA, NMI, and ARI values of 0.9, 0.904, 0.935, respectively. For the beta cell type with the biggest portion (47%), GraphSCC can correctly assign 98% cells. In contrast, the second best method, Seurat3.0 can correctly assign only 35% cells. Other methods make an accuracy of 34-67% on the cell type. Two major sources of wrong assignments in GraphSCC are the separation of the ductal cells into two clusters and merging of the gamma cells with another cluster. The separation of ductal cells was also seen in the SIMLR method, and the merging of the gamma cells was seen in the Seurat3.0. These similar mistakes may come from the difficulty to cluster these cell types.

**Fig. 5.**
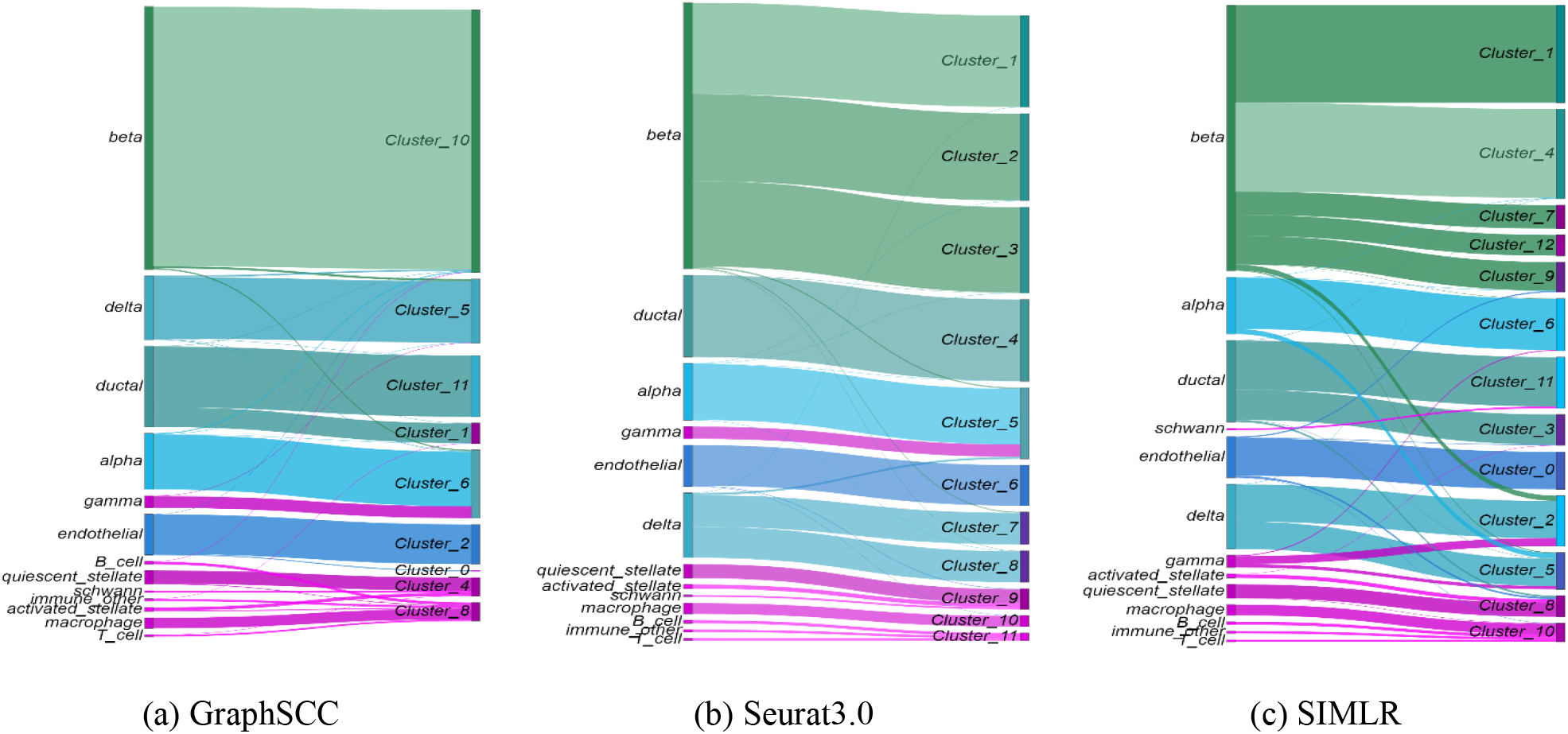
A Sankey river plot shows the match between clustering results and the actual labels on the Baron Mouse for three methods.

### C. Selecting Marker Genes and Functional Analysis

In this section, we selected the most differentially expressed genes for each cluster grouped by GraphSCC as the maker genes and examined if they are restricted to specific cell-type by public databases of cell-type markers. We selected PanglaoDB database[44] as the public dataset, which provides different marker genes for the same population. For example, PanglaoDB database provides 110 markers for B cells. Besides, we conducted an enrichment (GSE) analysis based on the selected marker gene set by GraphSCC.

As recommended in [45], we identify differentially expressed genes of each cluster relative to all other clusters using the FindAllMarkers function implemented in Seurat 3.0. Concretely, we selected sets of top 10 differentially expressed (DE) cluster-specific genes for each cluster. After we get the cluster-specific genes, we examined whether they are the marker genes by searching in PanglaoDB database (we only confirmed the cell types found in PanglaoDB database). As seen in Figure 6., the orange line showed the proportion of the marker genes in the predicted top 10 cluster-specific genes that can be verified in PanglaoDB database. The results showed that an average of 46% of predicted cluster-specific genes could be found in the PanglaoDB database, which was higher than the proportion of the total marker genes in whole genes. We also used the above approach to selected the top 10 cluster-specific genes based on the gold clusters (clusters consist of true cell type) shown in the green line. We can see, the number of marker genes we predicted is near to gold clusters, which indicates the accuracy of our clusters is high. Similar results were achieved for the top 5 and 20 cluster-specific genes predicted by GraphSCC (detailed information in **Figure S4**). Moreover, the detailed cluster-specific genes for each cell type were listed in the supplement **top_marker_genes**.**xls**. For cell types that can’t find in PanglaoDB database, our predicted genes for each cell type can be used as marker genes for further research. On the other hand, we performed functional analysis on the dataset Zeisel. As shown in Fig 7. (A), we selected the top 5 most differentially genes for each cell and found that the selected genes were highly differentially expressed in GraphSCC predicted cell type. Then, we performed biological pathway enrichment analysis based on the differentially expressed genes via the CompareCluster function implemented in clusterProfiler [46] R package along with default parameters. As shown in Fig 7. (B), each predicted cell type had its highly enriched pathway. For example, The literature [47] findings suggested that astrocytes in vitro may initially deploy cell-type-specific pat-terns of mRNA regulatory responses to glucocorticoids and subsequently activate additional cell type-independent responses.

**Figure 6.**
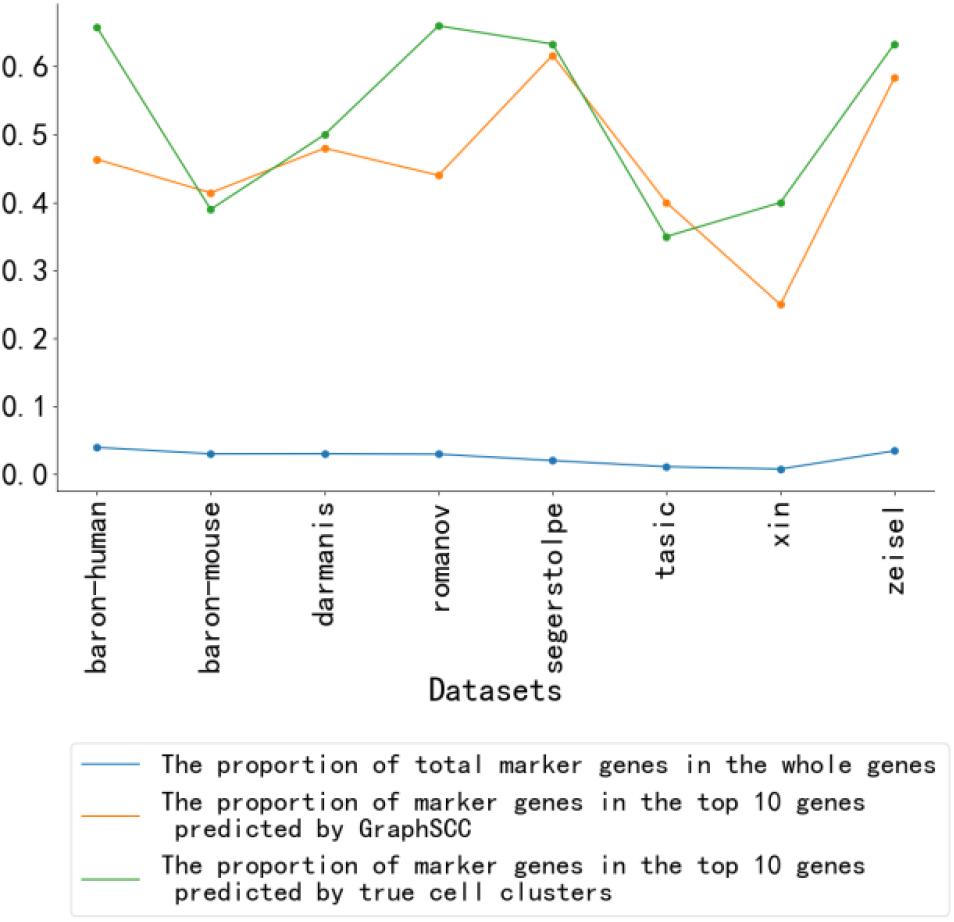
Line plots show that the proportion of real marker genes in top10 genes predicted by GraphSCC and true cell clusters. And the proportion of the sum of each cell’s marker genes (obtained from the PanglaoDB database) in the whole genes.

**Fig 7.**
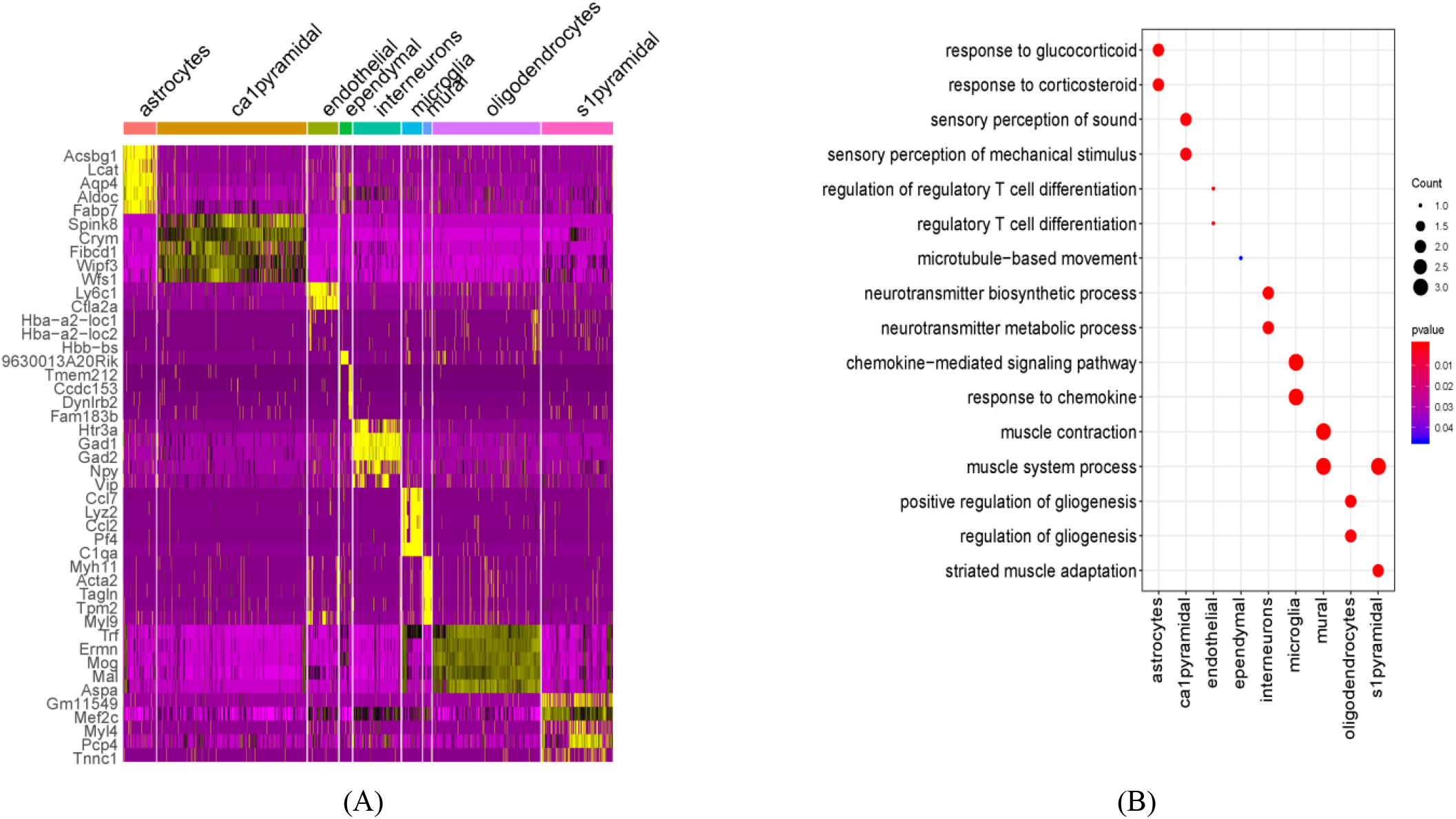
Zeisel dataset analysis based on GraphSCC. (A) Heatmap of DEGs (logFC>0.25) in each cell cluster. (B) enrichment pathways in each cell type using the top 5 differentially expressed genes.

### D. Contribution of Components to the Clustering

In this section, we investigated the contributions of components for the clustering performance of GraphSCC by conducting ablation studies on real datasets. As shown in TABLE II, the removal of the GCN module caused generally the largest drop in the performance with 7.8%, 4.8%, and 10.5% decreases in CA, NMI, and ARI, respectively. The changes indicate the importance of catching structural relations between cells. The removal of the residual connection (GraphSCC-Res) caused the 2nd biggest decreases in CA and NMI, while the biggest decrease in ARI. The residual connection is a good way to reduce the drawback of GCN to produce similar representations between nodes (cells), as indicated in the previous study [39]. The residual connection and more depth layers of GCN helped GraphSCC achieve better performance (**Figure S5**). We also clustered the cells based on learned distributions *Q* and *P* (denoted as GraphSCC (Q) and GraphSCC (P)), and they both cause a decrease in performances relative to GraphSCC that is based on distribution *Z*. In summary, the cooperation of the modules enables a better clustering of the scRNA-seq data.

**TABLE II.** ABLATION RESULTS ON REAL DATASETS

| Ablation tests | CA | NMI | ARI |
| --- | --- | --- | --- |
| GraphSCC-GCN | 0.72 | 0.743 | 0.639 |
| GraphSCC-Res | 0.768 | 0.736 | 0.672 |
| GraphSCC (Q) | 0.767 | 0.774 | 0.70 |
| GraphSCC (P) | 0.772 | 0.777 | 0.707 |
| GraphSCC | **0.798** | **0.791** | **0.744** |

## IV. Discussion and conclusion

This paper presents a structural deep clustering model GraphSCC consisting of GCN, DAE, and DSM modules. GraphSCC can effectively capture the relations between cells and the characteristics of data by learning representations using the GCN and DAE modules, respectively. Furthermore, DSM was applied to cluster cells based on representations by iteratively optimizing the clustering objective function in an unsupervised manner. We have demonstrated that the clustering performance of GraphSCC outperformed the competing methods on both simulated and real datasets. Furthermore, GraphSCC provided representations for better intra-cluster compactness and inter-cluster separability in the 2D visualization.

scRNA-seq is a revolutionary tool in biomedical research. Recently, many studies had been conducted based on scRNAseq technique. However, before we fully reap the benefit of scRNA-seq, many challenges must be overcome. Clustering cells into biologically meaningful groups is the critical step in scRNA-seq analyses. Through comprehensive evaluations with competing methods on real and simulated datasets, we have shown that GraphSCC offers stable clustering results based on scRNA-seq data. We believe that GrahpSCC will be a valuable tool for catching cellular heterogeneity. In the future, for better modeling the distribution of scRNA-seq data, we will integrate an imputation mechanism into GraphSCC. We will also apply graph transformer models and attention mechanisms to make scRNA-seq analyses more explainable.

## ACKNOWLEDGMENT

This study has been supported by the National Natural Science Foundation of China (61772566, 81801132, and U1611261), Guangdong Key Field R&D Plan (2019B020228001 and 2018B010109006) and Introducing Innovative and Entrepreneurial Teams (2016ZT06D211).

## AVAILABILITY OF DATA AND MATERIALS

The datasets we used in this study can be available at https://hemberg-lab.github.io/scRNA.seq.datasets/; All source code and datasets used in our experiments have been deposited at https://github.com/biomed-AI/GraphSCC.

## Notes

### Competing Interest Statement

The authors have declared no competing interest.

